# Rural selection drives the evolution of an urban-rural cline in coat color in gray squirrels

**DOI:** 10.1101/2023.02.02.526896

**Authors:** Bradley J. Cosentino, John P. Vanek, James P. Gibbs

## Abstract

Phenotypic differences between urban and rural populations are well-documented, but the evolutionary processes driving trait variation along urbanization gradients are often unclear. We combined spatial data on abundance, trait variation, and measurements of fitness to understand cline structure and test for natural selection on heritable coat color morphs (melanic, gray) of eastern gray squirrels (*Sciurus carolinensis*) along an urbanization gradient. Population surveys using remote cameras and visual counts at 76 sites along the urbanization gradient revealed a significant cline in melanism, decreasing from 48% in the city center to <5% in rural woodlands. Among 76 translocated squirrels to test for phenotypic selection, survival was lower for the melanic than gray morph in rural woodlands, whereas there was no difference in survival between color morphs in the city. These results suggest the urban-rural cline in melanism is explained by natural selection favoring the gray morph in rural woodlands combined with relaxed selection in the city. Our study illustrates how trait variation between urban and rural populations can emerge from selection primarily in rural populations rather than adaptation to novel features of the urban environment.

## 1. Introduction

In an era of unprecedented global change, how urbanization affects the evolution of life is a frontier in biology (Donihue and Lambert 2015, Johnson and Munshi-South 2017, Rivkin 2019). Urban areas are the fastest growing ecosystem on Earth (United Nations 2018), transforming land cover, climate, and hydrology, and creating novel interactions between people and wildlife (Grimm et al. 2008, Soulsberry and White 2015). These environmental changes can drive the evolution of novel adaptations within populations (Lambert et al. 2021). Yet, whereas phenotypic differences between urban and rural populations are well-documented (McDonnell and Hahs 2015, Alberti et al. 2017), the evolutionary processes controlling trait variation along urbanization gradients are less clear.

Phenotypic variation between urban and rural areas is not sufficient evidence of adaptive urban evolution. Phenotypic divergence can be due to plasticity rather than genetic variation, such as birds adjusting song pitch to high frequencies in response to low-frequency city noise (e.g., Slabbekoorn and Peet 2003). Demonstrating urban adaptation first requires showing trait variation is at least partially heritable. Genetic divergence between urban and rural populations provides further evidence of adaptation but is not entirely sufficient. Environmental differences between urban and rural areas may cause divergent natural selection on heritable traits, but phenotypic and genetic divergence can also result from genetic drift due to founder events, or low effective population size in combination with limited gene flow along the urbanization gradient (Vasemägi 2006). For example, colonization of cities by European blackbirds (*Turdus merula*) resulted in reduced genetic variation in city populations and divergence in migratory behavior between urban and rural areas (Evans et al. 2009, 2011). Ultimately, urban adaptation requires heritable trait variation that leads to differences in fitness among individuals. Despite a proliferation of case studies of urban-rural trait variation, many including patterns consistent with urban adaptation (Winchell et al. 2022), few studies examine the fitness consequences of trait variation (Lambert et al. 2021).

Fitness consequences of heritable trait variation can provide much needed insight into alternative hypotheses on the spatial context of adaptive and non-adaptive mechanisms generating urban-rural clines. Smooth clines along urbanization gradients can result from spatially divergent natural selection in the presence of gene flow (Takahashi 2015). In the case of discrete polymorphisms, divergent natural selection favors different morphs in urban and rural environments, generating a pattern of morph-by-environment interaction for fitness (Fig. 1*a*). Alternatively, urban-rural clines can result from a combination of selection and genetic drift across the urbanization gradient. For example, selection may favor one morph in the city, whereas morph frequencies are controlled primarily by drift in the rural environment (i.e., urban adaptation; Fig. 1*b*). Under this hypothesis, the morph favored by selection in the city should rarely reach high frequency in rural populations due to drift alone, and on average the frequency of the morph favored in the city should be negatively related to distance from the city (Fig. 1*b*). Urban-rural clines can also be generated by the reverse hypothesis of selection operating primarily in the rural environment with drift controlling morph frequencies in the urban environment (i.e., rural adaptation; Fig. 1*c*). Finally, if genetic drift is the salient force in both urban and rural populations, we expect no differences in fitness between morphs in either environment, with no consistent urban-rural clines (Fig. 1*d*).

**Figure 1.**
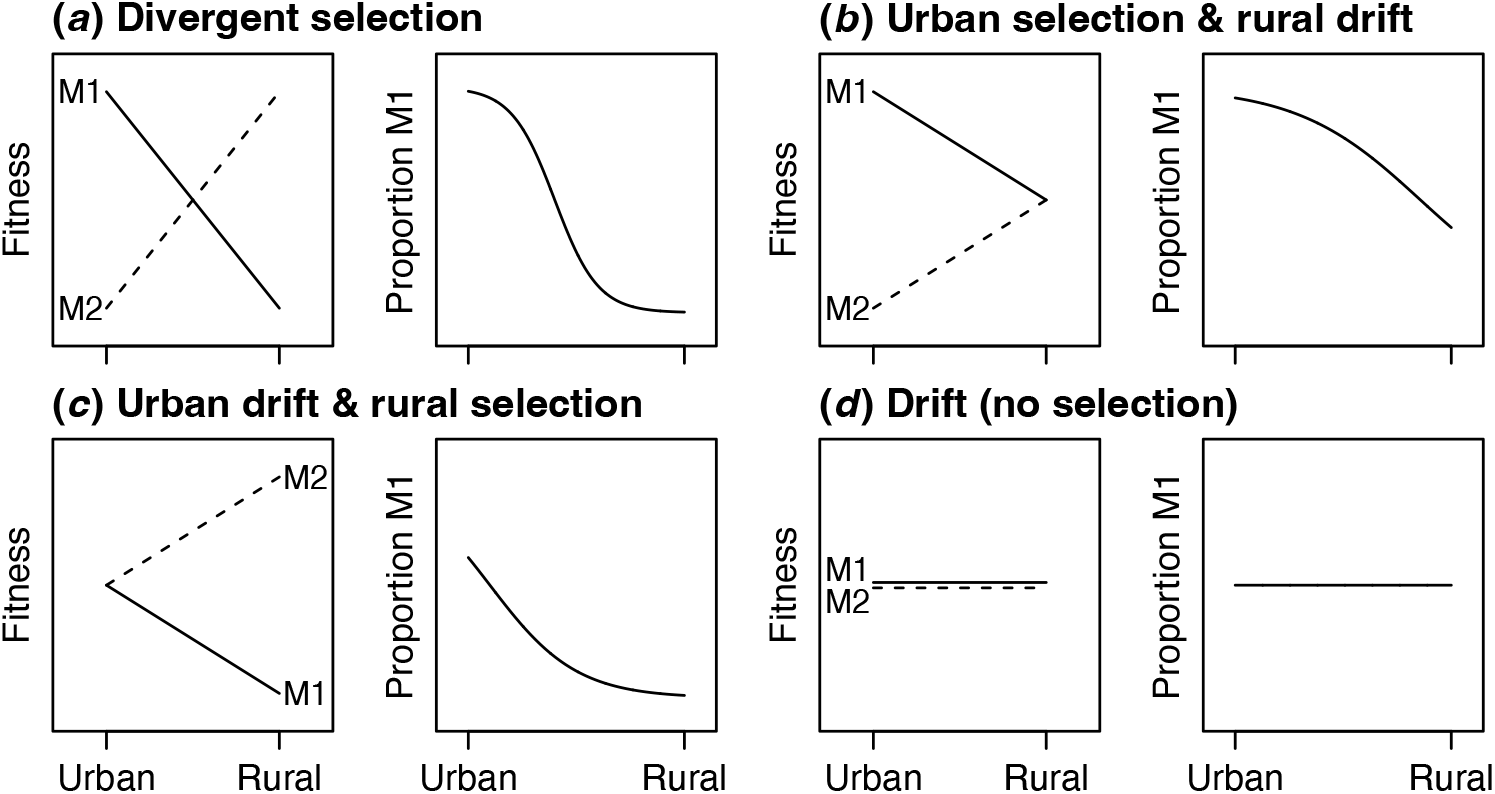
Four hypotheses (*a-d*) for spatial variation in the effects of selection and drift on a heritable, discrete polymorphism along urbanization gradients. Predicted relative fitness for two morphs (M1 and M2) in urban and rural environments and mean expectation for clinal variation in the proportion of morph M1 along urbanization gradients are shown for each hypothesis.

Our goal was to better understand the spatial context of adaptive and non-adaptive mechanisms of urban evolution. We used coat color in eastern gray squirrels (*Sciurus carolinensis*) as a model system. *Sciurus carolinensis* has two common color morphs – gray and melanic – controlled by the melanocortin-1-receptor gene (*Mc1R*; McRobie et al. 2009). Melanism in *S. carolinensis* is caused by a 24-bp deletion allele that is incompletely dominant, with heterozygotes having a brownish-black dorsum and brown underbelly (McRobie et al. 2009). Homozygotes for the deletion allele are black, whereas homozygotes for the recessive wild-type allele are gray. Melanics were the prevailing color morph in the northeastern United States prior to massive deforestation during European colonialism in the 18^th^ and 19^th^ centuries (e.g., Allen 1943, Schorger 1949, Robertson 1973), but today melanics tend to be rare in rural forests and more common in urban areas. Urban-rural clines in melanism occur across multiple cities at varying degrees of strength, and clines are absent in some cities (Cosentino and Gibbs 2022). These urban-rural clines are most evident in regions with relatively cold winter temperatures, where the deletion allele for melanism is likely maintained by selection for greater potential for non-shivering thermogenesis (Ducharme et al. 1989).

We combined two commonly used methods (see Linnen and Hoekstra 2009) to test whether urbanization causes natural selection on *S. carolinensis* coat color in Syracuse, New York, USA and surrounding rural areas. First, we characterized spatial variation in the prevalence of melanism along the urbanization gradient. Previous studies documenting an urban-rural cline in this area were based on observations of squirrels incidentally encountered via community science, which likely have spatial biases in sampling effort and detection of rare color morphs (Cosentino and Gibbs 2022). Here we used standardized surveys via remote cameras and visual counts and a novel hierarchical model that accounts for imperfect detection probability to test for an urban-rural cline. Second, we experimentally translocated squirrels of each color morph from urban to both urban and rural areas and compared survival between morphs in each environment to test for differential fitness. Previous work suggested an urban-rural cline in melanism may be related to two ecological causes of selection (Gibbs et al. 2019, Bryan 2020): 1) selection *against* melanics in rural forests because they are more conspicuous to predators and human hunters than the gray morph, and 2) selection *for* melanics in cities because they are more visible to drivers than the gray morph on roads and less likely to be struck by vehicles. Densities of *S. carolinensis* are significantly greater in urban than rural areas (Merrick et al. 2016), potentially due to relaxed predation pressure in cities relative to rural areas where *S. carolinensis* density is limited by predators (Havera and Nixon 1980, Bowers and Breland 1996). The predominant mortality source for urban tree squirrels is thought to be collisions with vehicles (McCleery et al. 2008). Thus, we predicted survival would be greater for the gray than melanic morph in rural forests where predation is the main driver of mortality, whereas survival would be greater for the melanic than gray morph in the urban environment where vehicular collision is the main driver of mortality.

## 2. Methods

### Study area

Syracuse (43.04068° N, -76.14373° W) is a medium-sized city (ca. 150,000 people) with a land area of 65 km^2^ and population density of ca. 2,300 persons/km^2^ (U.S. Census Bureau 2022). Mean tree cover is 27%, ranging from 4.5% at the city core to 47% at its periphery (Nowak et al. 2001), with rural areas outside the city consisting of a mosaic of forest and agriculture (Fig. 2).

**Figure 2.**
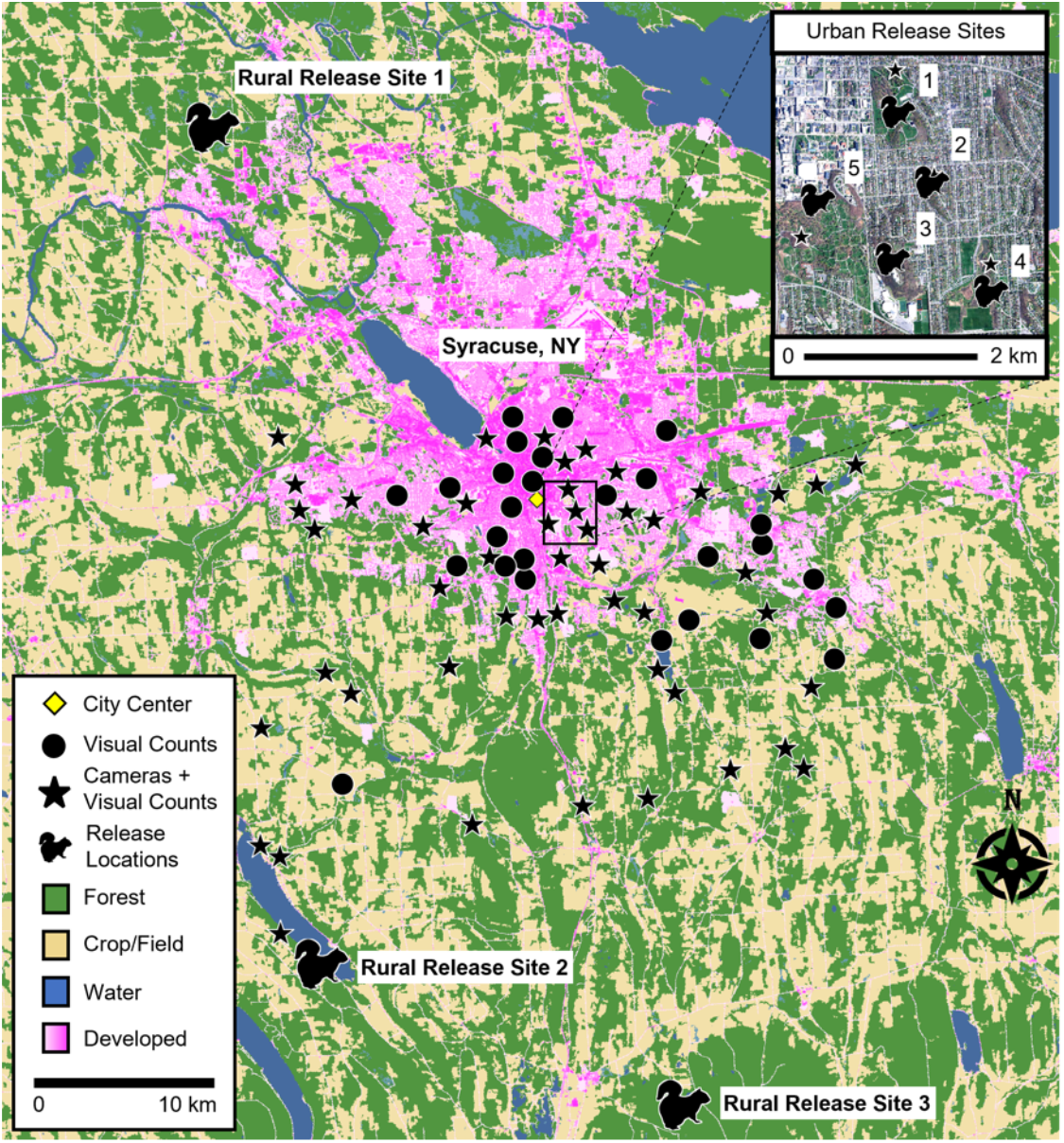
Land cover map of Syracuse, New York and surrounding rural areas. Sites with camera traps and visual counts (stars; *n* = 49) of eastern gray squirrels (*Sciurus carolinensis*) and sites with visual counts only (circles, *n* = 27) were distributed across the urbanization gradient. Rural release sites (*n* = 3) and urban release sites (*n* = 5; inset) for the translocation experiment are depicted by squirrel silhouettes. Land cover data modified from the National Land Cover Database (Yang et al. 2018). Digital orthoimagery from New York State Office of Information Technology Services GIS Program Office (2022a).

### Camera trapping and visual surveys

We estimated the relative abundance of each *S. carolinensis* color morph at 76 sites along a gradient of urbanization (Fig. 2). Squirrel occurrence is limited by tree cover, so we focused surveys on areas within urban greenspaces and rural woodlots with mature deciduous trees (Allen 1982, Parker and Nilon 2008). Sites surveyed were constrained to a sampling area that extended from the city center to rural woodlands 24 km south of the city. Site locations were chosen to represent a gradient in distance from the city center (range = 0.9–23.8 km, median = 6.8 km) with sites separated by at least 1 km.

We surveyed sites using a combination of remote cameras (i.e., camera trapping; Burton et al. 2015) and visual surveys. Camera trapping was conducted at 49 sites between 16 September 2021 and 13 October 2022. A single camera trap (Browning Strikeforce Pro XD) was deployed at each site between September 2021 and March 2022. Each camera was fastened to a tree 30-50 cm off the ground, generally facing north and aimed toward the ground with no obstructions within 2 m. All cameras were removed by October 2022, and cameras were active for 46-379 days (median = 263 days). Images from cameras were viewed by a single observer and classified to generate daily detection histories denoting whether each color morph was detected at each site on a given day.

We conducted visual surveys for squirrels at all camera sites and an additional 27 sites without cameras between 12 October 2021 and 18 October 2022 (*n* = 258 visual surveys). These additional sites were included to fill sampling gaps where permanent camera traps could not be placed. During each survey, a single observer recorded all visible squirrels of each color morph over a 3-min period. Each site was surveyed visually 1-7 times during the study (median = 4). Most sites (*n* = 57) received >1 visual survey across multiple days to facilitate abundance estimation with a hierarchical model that included an explicit model for the detection process. Median time between visual surveys was 65 days (range 10–197).

### Statistical analysis for cline estimation

We used an integrated hierarchical model (Kéry and Royle 2021) combining camera detection data with visual counts to estimate squirrel abundance and the proportion melanic at each site. The model is hierarchical in that it includes 1) an ecological process model to describe variation in squirrel abundance and the proportion melanic among sites and 2) an observation model to describe variation in detection probability of squirrels during surveys due to measurement error. We used the model to estimate the latent total abundance of squirrels (*N_total_i_*), the latent abundance of each color morph (*N_ik_*, where *k* = 1 for the melanic morph and *k* = 2 for the gray morph), the expected proportion of melanic squirrels (*p_melanic_i_*), and individual detection probabilities (*p_ijk_*) based on repeated surveys *j* of color morph *k* at each location *i*.

We included two submodels to describe the observation process: 1) A binomial distribution describing the counts *y_ijk_* of each morph from visual surveys (Royle 2004): *y_ijk_|N_ik_ ∼ Binomial*(*N_ik_, p_ijk_*), and 2) a Bernoulli distribution describing the binary detection observations *y_ijk_* of each morph from cameras (Royle and Nichols 2003): *y_ijk_|N_ik_ ∼ Bernoulli*(*P*_ijk_*), where *P*_ijk_* = 1 - (1 - *p_ijk_*)*^Nik^*. Because squirrel activity is related to daily ambient temperature (e.g., Wassmer and Refinetti 2019), we modeled individual detection probability as a function of mean daily temperature from a nearby (8.5 km from city center) weather station with a complete record for the study period (Syracuse International Airport, NOAA Climate Data Online, https://www.ncei.noaa.gov/cdo-web/). The detection probability model was logit(*p*_ijk_) = *a*0*_k_* + *a*1*_k_*\*temperature*_ij_* + *a*2*_k_*\*temperature^2^*_ij_*, where for each color morph *k*, *a*0*_k_* represents the intercept, *a*1*_k_* represents a linear effect of daily temperature, and *a*2*_k_* represents a quadratic effect of temperature. We included a quadratic term for temperature because we expected peaks in squirrel activity in fall and spring at intermediate temperatures and lower squirrel activity in winter and summer at temperature extremes (Spritzer 2002; electronic supplementary material, Fig. S1). Regression coefficients for detection probability were estimated with a joint likelihood, with the coefficients assumed to be identical between the count and detection/nondetection datasets. This integrated model formulation allowed us to combine count and detection/nondetection data in a single model, which can improve accuracy and precision of parameter estimates (Kéry and Royle 2021).

We included two submodels to describe the ecological state process. First, we used a Poisson distribution to describe the latent total abundance of squirrels: *N_total_i_* ∼ *Poisson*(*λ_i_*), where λ*_i_* is the expected total abundance of squirrels at location *i*. We expected total squirrel abundance to be greatest in the city (Merrick et al. 2016), so we modeled total abundance as a function of distance to city center: log(*λ_i_*) = *ß*0 + *ß*1*distance*_i_*, where *ß*0 represents the intercept and *ß*1 represents the effect of distance to city center. We defined the city center as the centroid of the civil boundaries of the city of Syracuse (Fig. 2; New York State Office of Information Technology Services GIS Program Office 2022b). We chose distance to city center as a general spatial index of urbanization because Syracuse has a simple, circular shape, and distance to city center is strongly correlated with multiple metrics commonly used to measure urbanization (Moll et al. 2019; e.g., impervious cover within 1 km: *r* = -0.70; road length within 1 km: *r* = -0.72; settlement density within 1 km: *r* = -0.72), each of which can represent different causal effects on the prevalence of melanism. Second, we used a binomial distribution to describe the latent abundance of the melanic morph: *N_i1_* ∼ *Binomial*(*N_total_i_, p_melanic_i_*). To estimate the cline in melanism, we modeled the proportion of melanic squirrels as a function of distance to city center: logit(*p_melanic_i_*) = *ß*0 + *ß*1*distance*_i_*, where *ß*0 represents the intercept and *ß*1 represents the effect of distance to city center. The latent abundance of the gray morph was derived as *N_i2_* = *N_total_i_ – N_i1_*.

We used JAGS 4.3.0 (Plummer 2017) and the R package *jagsUI* (Kellner and Meredith 2021) in R (Version 4.0.2; R Core Team 2020) to fit the integrated model with a Bayesian approach and Markov chain Monte Carlo. Covariates were standardized (mean = 0, SD = 1) before fitting the model. We fit the model with noninformative normal prior distributions for regression coefficients (Gelman and Hill 2007). We ran three chains with 12,000 iterations, discarding the first 2000 iterations as burn-in and thinning the remaining iterations by 10. The resulting 3000 iterations were used to describe the posterior distribution for each parameter. We confirmed convergence of parameter estimates with the Gelman-Rubin statistic (R-hat < 1.1; Gelman and Hill 2007).

We used a model of abundance that assumes closure to births, deaths, immigration, and emigration within sites during the survey period because our focus was on spatial variation in abundance of color morphs along the urbanization gradient. The closure assumption is unlikely to be met because *S. carolinensis* has an extended reproductive period, and juveniles can be found year-round (Kirkpatrick and Hoffman 1960; Steele and Koprowski 2003). When closure is not met, the abundance parameter (*λ_i_*) should be interpreted to represent the number of squirrels associated with a given site during the survey period rather than the number of permanent residents at the site (Kéry and Royle 2021). To evaluate whether our results were sensitive to the closure assumption, we fit our model using the same parameterization but with a subset of data during a period when closure was more likely met (16 September 2021 - 15 March 2022).

### Translocation experiment

To test for phenotypic selection, we conducted a translocation experiment and compared squirrel survival between color morphs in urban and rural environments. Squirrels of both color morphs were translocated from urban greenspaces in Syracuse to both urban and rural release sites (Fig. 2; electronic supplementary material, Table S1). Although we preferred an experimental design with squirrels translocated from both urban and rural environments, the rarity of melanic squirrels in rural areas precluded a full reciprocal design, which ultimately prohibited us from testing for local adaptation (e.g., greater survival of each color morph in its home than away environment; Kawecki and Ebert 2004). However, our design allowed us to compare survival as an index of fitness between morphs within environments as a test of phenotypic selection (Linnen and Hoekstra 2009), which was our focus.

We measured survival via radiotelemetry. To do so, we captured and collared 76 squirrels in urban greenspaces between June 2021 and May 2022 using wire cage traps (Tomahawk Live Trap Co., Tomahawk, WI; Model 103) baited with peanut butter. Upon capture we coaxed squirrels into a cloth handling cone for processing and radio-collar attachment (Koprowski 2002). Squirrels were anesthetized using isoflurane delivered via a plastic nose cone to minimize squirrel stress and pain response and to maximize human safety (Parker et al. 2008). We recorded squirrel mass, color morph, sex, and reproductive status. We collected a 1×3-mm sliver of ear tissue for future genetic analyses and attached a metal ear tag (National Band Company, Newport KY; Style 1005-3) with a unique numeric code to the other ear. We attached VHF (“very high frequency”) radiotransmitters to squirrels via neck collars (Holohil Systems Ltd. Carp, ON; Model RI-2D 14 or 21g; with mortality switch option). We epoxied a metal temperature logger (Maxim Integrated DS1925L-F5, 3.49 g) to the end of collars to record temperature data. Weight of the radiotransmitter, collar, and temperature logger was <5% of the squirrel’s mass (mean = 2.7%, range = 2-4%). Pregnant and lactating females were excluded from the sample. Following radio-collar attachment and recovery from isoflurane, we transported each squirrel to a rural or urban release site by vehicle. To control for impacts of travel time, we drove all translocated squirrels for a similar amount of time before release (30 min). Although the process of translocation can be disruptive to an individual by removing it from its home range, any effect of translocation on survival was controlled because all individuals in our study were translocated.

Three rural release locations were 22-30 km from trapping locations and composed of deciduous and mixed deciduous forest. Five urban release locations were 0.7-6 km from trapping locations within Syracuse and consisted of a cluster of urban greenspaces with mixed deciduous forest. All urban and rural release locations had known populations of *S. carolinensis*. Release sites for each squirrel were selected systematically to maintain a balanced release of morph types across sites at locations with ample shelter (e.g. tree cavities) and food resources (mast-producing trees). In total, 18-20 squirrels of each morph were translocated to urban and rural environments (Table 1).

**Table 1.** Number of translocated eastern gray squirrels (*Sciurus carolinensis*) and their fates by color morph in each environment to which squirrels were translocated. Fates included confirmed & probable mortalities, possible mortalities (mortalities and collars that were transmitting a mortality signal, but we were unable to be locate), missing squirrels that we could no longer find using radiotelemetry, and squirrels with active collars at the end of the study (October 2022).

| Translocations and Fates | Color morph | Translocated environment |  |
| --- | --- | --- | --- |
|  |  | Urban | Rural |
| Translocated from urban environment | Melanic | 18 | 18 |
|  | Gray | 20 | 20 |
| Confirmed & probable mortalities | Melanic | 2 | 8 |
|  | Gray | 4 | 2 |
| Possible mortalities | Melanic | 10 | 8 |
|  | Gray | 6 | 12 |
| Missing | Melanic | 4 | 2 |
|  | Gray | 8 | 5 |
| Active | Melanic | 2 | 0 |
|  | Gray | 2 | 1 |

After release, squirrel survival was monitored weekly (Haughland et al. 2008; Millspaugh et al. 2012) until October 2022. At the end of the study, squirrel status was classified into one of four categories: 1) confirmed and probable mortalities, 2) possible mortalities, 3) missing squirrels that could no longer be found with radiotelemetry, or 4) active (Table 1). When we detected a squirrel’s radiotransmitter emitting a mortality signal, we attempted to locate the squirrel as soon as possible. Squirrels with a mortality signal were assigned to the “confirmed and probable” mortality category based on either the presence of a carcass (i.e., confirmed) or significant evidence of a mortality event in the absence of a carcass (e.g. deep, rounded bite marks on the collar assembly, large amounts of squirrel fur, or evidence the collar assembly was removed by a human; Adams et al. 2004, Tebo and Norton 2017, Wilson et al. 2019). When possible, we tried to ascertain the cause of mortality (McCleery et al. 2008). Possible mortalities were based on the presence of a transmitter with a mortality signal but without a carcass or other evidence of a depredation event, including transmitters broadcasting from a consistently immobile position that we were unable to retrieve (e.g., emanating from a treetop).

### Statistical analysis for survival estimation

We used a Cox proportional hazards model (Cox 1972; Pollock et al. 1989) to relate daily survival of translocated squirrels to color morph (gray, melanic) and translocation environment (urban, rural). Date of translocation was used as the time of entry to maximize the number of squirrels at risk at a given time for estimating daily survival (Karp and Gehr 2020). We used the inferred date of mortality as the exit date in the model for squirrels classified as confirmed and probable mortalities. Date of mortality was estimated as the median date between when the last active signal was received and the first instance a mortality signal was detected. For collars recovered with functioning temperature loggers (*n* = 21), we determined the date of mortality as the date when collar temperature remained at ambient temperature, no longer rising to core diurnal body temperature for *S. carolinensis* (37.9 °C; Pereira et al. 2002, Wassmer and Refinetti 2019). Squirrels not classified as mortalities contributed to the survival analysis via right censoring, as each had a known minimum number of days survived (Murray 2006).

We modeled the hazard ratio as a function of morph, environment, and the interaction between morph and environment using the R package *survival* (Therneau 2021). A likelihood ratio test was used to examine the significance of the interaction term, with two-sided Wald tests reported for each term in the most-supported model. We considered a model with additional covariates, including sex, body mass, collar size, and mean daily temperature during the first week after translocation (linear and quadratic effects), but we excluded these variables from the final model because a) none of the variables predicted the hazard ratio (electronic supplementary material, Table S2), b) data on sex was missing for two squirrels, which precluded use of those observations in a model with sex as a covariate, and c) the effects of morph, environment, and their interaction were consistent with or without the additional covariates in the model. No deviations from the assumption of proportional hazards were evident through analysis of the Schoenfeld residuals for morph (χ^2^ = 0.24, d.f. = 1, *P* = 0.62), environment (χ^2^ = 2.20, d.f. = 1, *P* = 0.14), the morph-environment interaction (χ^2^ = 0.01, d.f. = 1, *P* = 0.93), or the global model (χ^2^ = 4.06, d.f. = 3, *P* = 0.26). Kaplan-Meier estimates were used to quantify survival of each color morph over time, which were visualized with the *survminer* package (Kassambara et al. 2021). We also used log rank tests for pairwise comparisons of survival between morphs within each environment. Finally, to test whether the results were sensitive to our classification of mortalities, we refit the Cox proportional hazards model and log rank tests classifying both “confirmed and probable” and “possible” mortality categories as true mortalities.

## 3. Results

### Cline estimation

Across the 76 sites sampled, we detected the melanic morph at 40 sites (53%) and the gray morph at 64 sites (84%) on either point counts or camera imagery. We counted 237 squirrels across the 76 sites during 258 visual surveys, including 69 melanic morphs (0-8 per survey) and 168 gray morphs (range = 0-7 per survey). Among the 49 sites with cameras, the melanic morph was detected at 35 sites (71%), and the gray morph was detected at all sites (100%). The median number of days detected on camera at each site was 23 for the melanic morph (range: 0-321) and 111 for the gray morph (range: 5-351).

The proportion of melanic squirrels was negatively related to distance from the city center, declining from 48% at the city center to <5% in rural woodlands (Fig. 3; electronic supplementary material, Table S3). Total squirrel abundance was also negatively related to distance from city center (Fig. 4; electronic supplementary material, Table S3), but abundance of the melanic morph declined at a rate faster than abundance of the gray morph with increasing distance from the city center (Fig. 4). Detection probability of both morphs peaked at intermediate daily mean temperature, and detection probability was consistently greater for the gray than melanic morph (electronic supplementary material, Fig. S2). All model parameters converged (R-hat < 1.01), with results virtually identical when fitting the model with a subset of data when population closure was more likely met (electronic supplementary material, Table S4, Figs. S3-S5).

**Figure 3.**
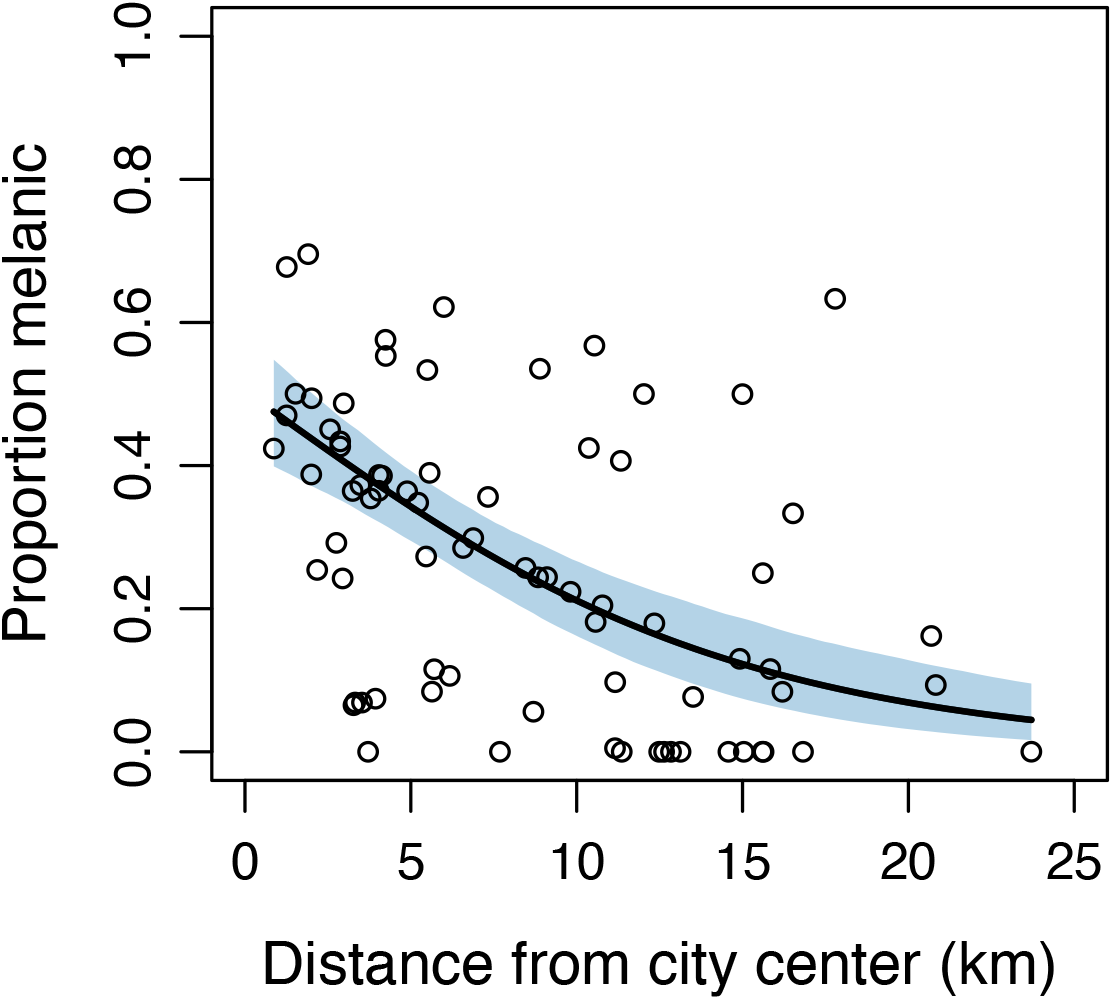
Relationship between the proportion of melanic eastern gray squirrels (*Sciurus carolinensis*) and distance to city center for Syracuse, New York based on a hierarchical model integrating camera trap and visual count observations of squirrels across *n* = 76 sites (Fig. 2). Solid line represents the predicted proportion of melanic squirrels from the model, and shaded area represents a 95% credible interval. Open circles represent estimates of the proportion of squirrels melanic at each site.

**Figure 4.**
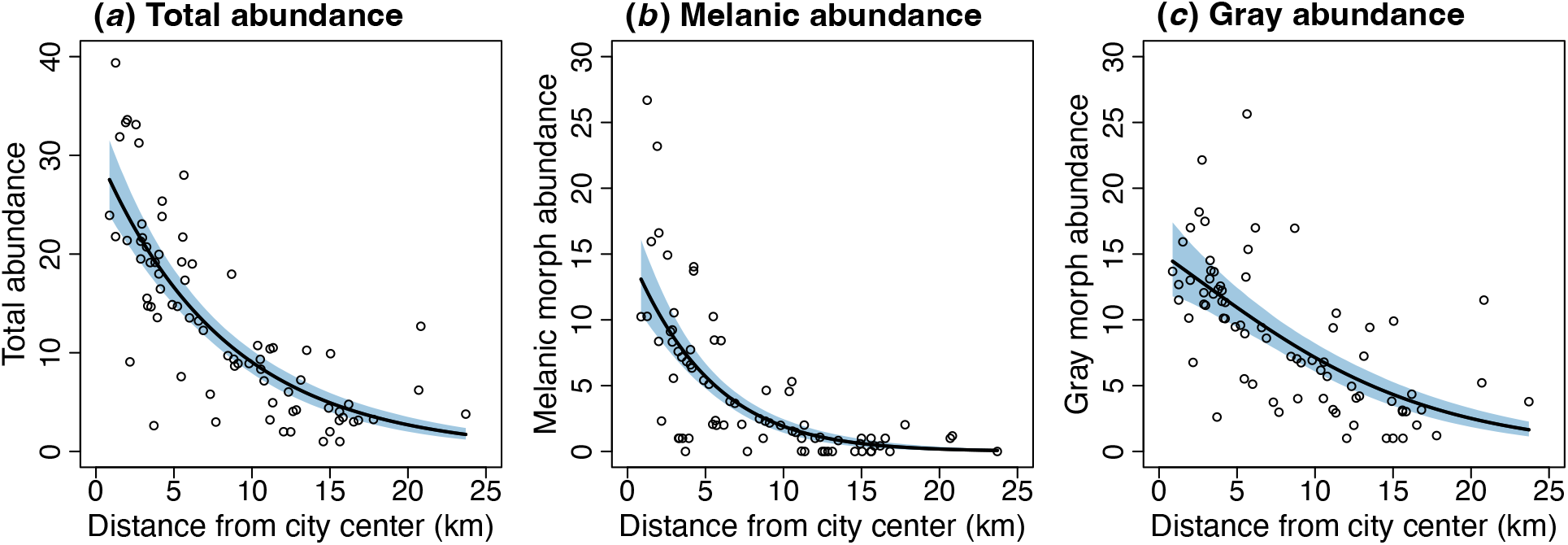
Relationship between the abundance of eastern gray squirrels (*Sciurus carolinensis*) and the distance to city center for Syracuse, New York, in total (*a*) and for each color morph (*b-c*), based on a hierarchical model integrating camera trap and visual count observations of squirrels across *n* = 76 sites (Fig. 2). Solid lines represent predicted abundance from the model, and shaded areas represent 95% credible intervals. Open circles represent estimates of abundance at each site.

### Survival estimation

Of the 76 squirrels collared and translocated, we documented 16 confirmed and probable mortalities by the end of the tracking period (Table 1). An additional 36 squirrels with mortality signals were classified as possible mortalities. Nineteen squirrels went missing during the study, and five squirrels were still active at the end of the study. The Cox proportional hazards model revealed an interaction effect between morph and environment on the mortality hazard ratio (Likelihood ratio test: χ^2^ = 5.70, d.f. = 1, *P* = 0.017). The model with the interaction term indicated significant hazard ratios for morph, environment, and the morph*environment interaction (Table 2). The melanic morph had eight times the mortality risk as the gray morph in the rural environment (hazard ratio for gray relative melanic in rural areas = 0.12, 95% CI: 0.03 –0.60), whereas there was no difference in the odds of mortality between melanic and gray morphs in the urban environment (electronic supplementary material, Fig. S6). Post-hoc log rank tests showed survival was lower for the melanic than gray morph in the rural environment (Fig. 5; χ^2^ = 7.2, d.f. = 1, *P* = 0.007), whereas there was no difference in survival between color morphs in the urban environment (Fig. 5; log rank test: χ^2^ = 0.4, d.f. = 1, *P* = 0.50).

**Figure 5.**
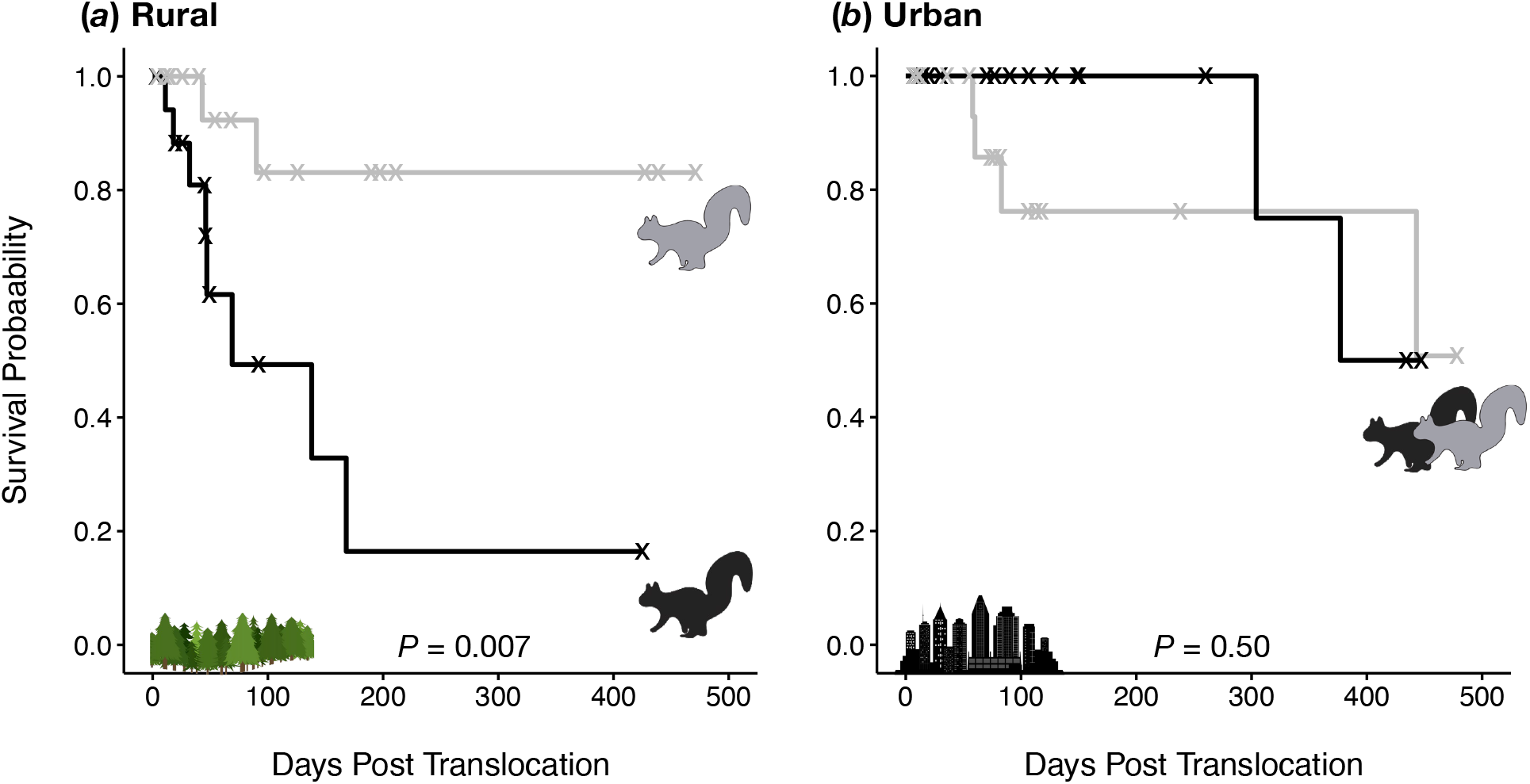
Survival curves for gray (solid gray line) and melanic (solid black line) color morphs of eastern gray squirrels (*Sciurus carolinensis*) translocated to rural (*a*) or urban (*b*) environments (see Table 1 for sample sizes). Each step down represents a confirmed or probable mortality (Table 1), and “X”s represent possible, missing, and active squirrels that were right-censored for the analysis. *P*-values indicate the effect of morph on survival within each environment based on log rank tests.

**Table 2.** Parameter estimates for Cox proportional hazards regression model for effects of color morph (melanic, gray) and environment (rural, urban) on daily survival of translocated eastern gray squirrels (*Sciurus carolinensis*). Reference categories were set as melanic for color morph and rural for environment.

| Explanatory variable | Log hazard ratio | SE | $z$ | $P$ |
| --- | --- | --- | --- | --- |
| Color morph | -2.09 | 0.80 | -2.61 | 0.009 |
| Environment | -2.12 | 0.80 | -2.64 | 0.008 |
| Color morph * Environment | 2.65 | 1.18 | 2.24 | 0.025 |

Results of the survival analysis were largely the same when including both the 16 confirmed and probable mortalities and 36 possible mortalities as true mortalities in the analysis. The Cox proportional hazard model showed an interaction effect between morph and environment (Likelihood ratio test: χ^2^ = 3.84, d.f. = 1, *P* = 0.05), and there was an additional effect of collar size on the mortality hazard ratio (electronic supplementary material, Table S5). Squirrels with the large collar had a greater hazard than squirrels with the small collar (electronic supplementary material, Fig. S7), but the interpretation of the morph*environment interaction remained the same. Survival was lower for the melanic than gray morph in the rural environment (electronic supplementary material, Fig. S8; log rank test: χ^2^ = 4.2, d.f. = 1, *P* = 0.04), whereas there was no difference in survival between melanic and gray morphs in the urban environment (log rank test: χ^2^ = 0.2, d.f. = 1, *P* = 0.60)

## 4. Discussion

Urbanization is hypothesized to foster adaptation to features novel to cities, but empirical studies differentiating between adaptive and non-adaptive drivers of urban evolution have been limited (Lambert et al. 2021). Our work suggests a combination of adaptive and non-adaptive processes can generate urban-rural clines in phenotypic traits. We found evidence of a strong urban-rural cline in melanism in *S. carolinensis* in our study area, with melanism increasing tenfold from the rural areas to the city center. Our translocation experiment provides insight into the evolutionary mechanisms generating this cline. We found evidence of natural selection on coat color in the rural environment, where squirrel survival was significantly greater for the gray than melanic morph. In contrast, there was no evidence of selection on coat color in the urban environment, where squirrel survival was similar between color morphs. These results suggest that adaptive mechanisms of urban evolution do not necessarily result from urban adaptation *per se* (i.e., adaptation in the city proper), and instead that urban-rural clines in heritable traits can emerge from selection in the rural environment alone.

Our finding of an urban-rural cline in melanism is consistent with previous studies on coat color in *S. carolinensis* (Gibbs et al. 2019, Lehtinen et al. 2020, Cosentino and Gibbs 2022) and more broadly with the body of evidence showing phenotypic divergence between urban and rural populations (McDonnell and Hahs 2015, Alberti et al. 2017). The cline unveiled here was significantly stronger than the cline predicted for Syracuse from community science observations (Cosentino and Gibbs 2022), underscoring the importance of conducting standardized surveys (documenting both presence and absence data) and estimating the relative abundance of trait variants with a model that accounts for imperfect detection probability. Notably we found that detection probability during surveys peaked at moderate temperatures typical of spring and fall seasons (electronic supplementary material, Figs. S2), but with detection probability greater for the gray than melanic morph at all but the coldest temperatures (electronic supplementary material, Fig. S2). The melanic morph is generally more visible to human observers than the gray morph (Gibbs et al. 2019, Bryan 2020), so the difference in detection probability between morphs is not likely explained by differences in crypsis. It is possible that melanic *S. carolinensis* individuals have lower activity levels than gray individuals, effectively decreasing the availability of melanics to be observed during surveys. Eumelanin production via the melanocortin system is often correlated with a range of physiological and behavioral traits (Roulin and Ducrest 2011), so further evaluation of behavioral differences between color morphs in *S. carolinensis* is needed. Regardless of the mechanisms explaining the difference in detection probability between morphs, our findings highlight the importance of considering intraspecific variation in detection probability when fitting hierarchical models of abundance.

Cities are commonly thought to be engines of evolution by creating novel ecological conditions that cause positive selection (Shochat et al. 2006). Indeed, urban evolution studies with strong evidence of adaptive trait variation (*sensu* Lambert et al. 2021) have all found novel selection pressures in the city. For example, Atlantic killifish (*Fundulus heteroclitus*) evolved tolerance to lethal pollutants in urban estuaries (Nacci et al. 2010), and urban water fleas (*Daphnia magna*) evolved a greater critical thermal maximum under urban heat stress (Brans et al. 2017). Fitness tradeoffs have also been documented between urban and rural populations. In the textbook case of peppered moths (*Biston betularia*), selection favored the dark morph in polluted cities with dark tree backgrounds and the light morph in unpolluted rural areas with light tree backgrounds (Cook 2003, Cook et al. 2012). Similarly, urban acorn ants (*Temnothorax curvispinosus*) evolved greater heat tolerance than rural ants in the warm urban environment, whereas urban ants had lower cold tolerance than rural ants in the cool rural environment (Martin et al. 2019). In this study, melanic and gray morphs of *S. carolinensis* experienced similar levels of mortality when translocated within the urban environment, suggesting that melanism has not adaptively evolved in response to novel selection pressures in the city. Instead, urban-rural clines in melanism appear to emerge primarily from selection for the gray morph in rural populations, as gray survival was substantially greater than melanic survival among individuals translocated to rural woodlands. Our results suggest that selection in rural areas is relaxed in the city, and the dominant evolutionary process controlling coat color in city populations is likely genetic drift.

Our findings can help explain why the prevalence of melanism often varies substantially between nearby cities. Cosentino and Gibbs (2022) found that urban-rural clines in melanism in *S. carolinensis* tend to evolve repeatedly among cities, but they are not universal. Urban-rural clines in melanism are rare in regions with warm winters, where the prevalence of melanism is likely limited due to thermal selection (Ducharme et al. 1989, Cosentino and Gibbs 2022). But even within regions with colder winter temperatures where the melanic morph is regionally common, the prevalence of melanism can vary widely among cities. For example, Lehtinen et al. (2020) found the prevalence of melanism ranged 0-96% among cities in Ohio, and we found melanism ranged 0-50% among cities in upstate New York (Cosentino and Gibbs 2022; B.J. Cosentino, unpublished data). Dramatic variation in the frequency of melanism among nearby cities may be the result of differential introduction history and consequent founder effects, as melanic squirrels have commonly been introduced to cities as novelty (Benson 2013). Limited evidence about temporal variation in melanism in cities also suggests drift is the primary driver of color morph frequencies within the urban core. For example, Lehtinen et al. (2020) found no consistent change in the prevalence of melanism over eight years of monitoring in Wooster, Ohio, consistent with the stochastic effects of drift on morph frequencies (although it is possible for equilibrium conditions to emerge from a balance of selection with gene flow or mutation). More studies are needed to examine the mechanisms driving variation in cline strength among cities. Moreover, although we documented a strong spatial cline in melanism along the urbanization gradient in Syracuse, there is considerable variation in the prevalence of melanism within any segment of the urban-rural gradient (Fig. 3). Landscape heterogeneity within both urban and rural areas could cause microgeographic variation in the strength of selection, drift, and gene flow. Mechanistically linking fine-scale trait variation to landscape heterogeneity should be a priority for future urban evolution research.

Relaxed selection can result from environmental change weakening a source of selection on trait variation, resulting in either a new vector of selective costs and benefits or neutral evolution of the trait variation in question (Lahti et al. 2009). Environmental change in cities has been hypothesized to cause relaxed selection in several ways, such as relaxation of food and water limitation (Shochat et al. 2006). For example, Rodewald et al. (2011) described a putative case of relaxed selection in northern cardinals (*Cardinalis cardinalis*), where food subsidies in urban areas break a positive relationship between plumage color and individual condition found in rural areas where food is limited. Although our study does not provide insight into the ecological causes of selection, one hypothesis is that selection via predation and hunting favoring the gray morph in rural woodlands is relaxed in cities. *Sciurus carolinensis* has a variety of avian and mammalian predators and have been hunted by people for centuries (Benson 2013). Our previous work showed the gray morph is more cryptic against tree backgrounds than the melanic morph in regrown, secondary forests that dominate the landscape in rural New York (Gibbs et al. 2019, Bryan 2020), potentially leading to lower levels of predation on the gray morph. Selection via predation is likely weaker in cities than rural secondary forests. Hunting is nearly universally banned in cities, and predation rates are often lower in cities than rural areas as food subsidies (e.g., bird seed, pet food, garbage) redirect foraging of predators away from some prey (Fischer et al. 2012). Multiple lines of evidence suggest predation rates on squirrels are lower in urban than rural areas. For example, experiments on giving-up-densities and food supplementation have indicated that rural *S. carolinensis* populations are predator-limited, whereas urban populations are food-limited (Havera and Nixon 1980, Bowers and Breland 1996). Moreover, flight initiation distances from an approaching predator are lowest where human exposure is greatest (Engelhardt and Weladii 2011), and a study of eastern fox squirrel (*S. niger*) mortality showed >60% of fatalities were caused by predation at a rural site compared to <5% at an urban site (McCleery et al. 2008). Ultimately experimental study of attack rates on color morphs of *S. carolinensis* in urban and rural areas is needed to clarify whether predation is a cause of selection on coat color in rural populations (i.e., greater attack rates on the melanic than gray morph) and to test whether selection via predation is relaxed in cities. Given the correlation of melanism with other physiological and behavioral traits (Roulin and Ducrest 2011), future studies should also clarify the degree to which selection on melanism is direct (e.g., crypsis) versus indirect via correlations with other traits or early life conditions.

Similar survival between color morphs in the urban environment was counter to our prediction. The melanic morph is more visible to humans than the gray morph on roads (Bryan 2020), and community science data suggested the melanic morph was underrepresented among road killed squirrels in our study area (Gibbs et al. 2019), so we expected selection for the melanic morph in the city. The gray morph may be more susceptible to vehicular collisions without this source of mortality contributing to the urban-rural cline in melanism. For example, fitness costs of road mortality for the gray morph in urban areas (where traffic volume is greatest) could be balanced by other sources of fitness costs for the melanic morph (e.g., greater visibility to some urban predators than the gray morph), leading to no net difference in fitness between morphs in city populations.

Our translocation experiment was limited in multiple ways. First, because all squirrels in the survival analysis were translocated, we were unable to directly test if there was a significant effect of translocation on survival (e.g., greater survival of squirrels in their home range than a translocated environment). However, Koprowski et al. (2016) reports annual survival in *S. carolinensis* is often >0.50, which is consistent with annual survival estimates for all translocated squirrels in our study except the melanic morph in rural areas (Fig. 5). Second, we were rarely able to confirm the cause of mortality when retrieving radiotransmitters, and therefore unable to examine variation in the causes of mortality along the urbanization gradient. Nearly half of the translocated squirrels (36 of 76) had ambiguous fates despite an active mortality signal. These 36 squirrels were recorded as “possible” mortalities and included 14 cases where the radiotransmitter was unretrievable (e.g., at the top of a tree). Of the 22 possible mortalities where we retrieved the radiotransmitter, we found no evidence of physical damage. Some of these possible mortalities were likely due to slipped collars and did not represent true mortalities, which is why we refrained from including possible mortalities in our main survival analysis. When we conducted the survival analysis with these possible mortalities classified as true mortalities, the hazard of mortality was greater for squirrels with large than small collars, supporting the idea that possible mortalities are sometimes slipped collars (more likely for large collars). Nonetheless, we found a significant morph*environment interaction and the same pattern of differential survival between morphs in each environment regardless of whether possible mortalities were included as true mortalities in the survival analysis. Third, our study focused solely on the vital rate survival as a component of fitness. We lack data on differential reproductive success between morphs, although our focus on adult survival was important given that the best short-term proxies of lifetime fitness are those measured at the recruitment stage (Alif et al. 2022).

In conclusion, our study highlights how the effects of urbanization on trait variation can reach beyond the city boundaries, driving adaptive evolution in nonurban areas. The rise of cities was accompanied by conversion of adjacent rural lands – often distant from urban centers – to agriculture and other land uses for resource extraction to support urban populations (Cronon 1991). European settlements dramatically reshaped the landscape in the northeastern United States, converting extensive old growth forests to secondary forests that have regrown over the last 200 years (Foster et al. 2010, Thompson et al. 2013). It was this period of time during which melanism in *S. carolinensis* began to decline in rural forests (e.g., Allen 1943, Schorger 1949, Robertson 1973), reinforcing the idea that “urban evolution” not only encompasses the effects of contemporary selective regimes on trait variation in urban centers, but also the historical evolution of trait variation associated with landscape change beyond the city boundaries. Our study underscores the importance of comparing fitness between trait variants in both urban and rural populations to provide insight into the spatial context of adaptive and nonadaptive evolutionary processes along urbanization gradients.

## Supporting information

Electronic Supplementary Material

## Acknowledgements

This research was funded by the U.S. National Science Foundation (DEB-2018140, DEB-2018249). Field assistance was provided by Regina Hashim, Thomas Klein, Jessica Proctor, Richard Rich, Mark Suchewski, and Joelee Tooley. Ben Augustine provided feedback on hierarchical models. Access to field sites was provided by the New York Department of Environmental Conservation, City of Syracuse, and select private and municipal landowners.

