## Supplementary material for "Rural selection drives the evolution of an urban-rural cline in coat color in gray squirrels": Electronic Supplementary Material

B.J. Cosentino, J.P. Vanek, and J.P. Gibbs

**Guide to Supplementary Figures and Tables**

**Figure S1.** Relationship of mean daily temperature to survey date during the period when camera trapping and visual surveys were conducted for eastern gray squirrels (*Sciurus carolinensis*).

**Figure S2.** Relationship of individual detection probability of melanic and gray morphs of eastern gray squirrels (*Sciurus carolinensis*) to daily mean temperature based on a hierarchical model integrating camera trap and point count observations across N = 76 sites.

**Figure S3.** Relationship of the proportion of melanic eastern gray squirrels (*Sciurus carolinensis*) to the distance to city center for Syracuse, NY based on a hierarchical model integrating a subset of camera trap and point count observations of squirrels across N = 50 sites (16 September 2021 – 15 March 2022).

**Figure S4.** Relationship of the abundance of eastern gray squirrels (*Sciurus carolinensis*) in total and for each color morph to the distance to city center for Syracuse, NY based on a hierarchical model integrating a subset camera trap and point count observations of squirrels across N = 50 sites (16 September 2021 – 15 March 2022).

**Figure S5.** Relationship of individual detection probability of melanic and gray morphs of eastern gray squirrels (*Sciurus carolinensis*) to daily mean temperature based on a hierarchical model integrating a subset of camera trap and point count observations across N = 50 sites (16 September 2021 – 15 March 2022).

**Figure S6.** Log hazard ratios predicted from the Cox proportional hazards regression model including effects of morph, environment, and morph*environment interaction (Table 2).

**Figure S7.** Log hazard ratios predicted from the Cox proportional hazards regression model when including confirmed, probable, and possible mortalities as true mortalities.

**Figure S8.** Survival curves for gray (solid gray line) and melanic (dotted black line) color morphs of eastern gray squirrels (*Sciurus carolinensis*) translocated to rural (A) or urban (B) environments when including confirmed, probable, and possible mortalities as true mortalities.

**Table S1**. Locations of sites to which eastern gray squirrels (*Sciurus carolinensis*) were translocated for survival estimation in urban and rural environments.

**Table S2.** Parameter estimates for Cox proportional hazards regression model for effects of color morph (melanic, gray), environment (rural, urban), sex, body mass, collar size, and mean daily temperature during the first week of translocation on daily survival of translocated eastern gray squirrels (*Sciurus carolinensis*).

**Table S3.** Parameter estimates and 95% Bayesian credible intervals (CI) for models of abundance, proportion melanic, and detection probability of eastern gray squirrels (*Sciurus carolinensis*) based on a hierarchical model integrating camera trap and point count observations across N = 76 sites.

**Table S4.** Parameter estimates and 95% Bayesian credible intervals (CI) for models of abundance, proportion melanic, and detection probability of eastern gray squirrels (*Sciurus carolinensis*) based on a hierarchical model integrating a subset of camera trap and point count observations across N = 50 sites (September 2021 – 15 March 2022).

**Table S5.** Parameter estimates for Cox proportional hazards regression model for effects of color morph (melanic, gray), environment (rural, urban), and collar size (14 g vs. 21 g) on daily survival of translocated eastern gray squirrels (*Sciurus carolinensis*) when including confirmed, probable, and possible mortalities as true mortalities.

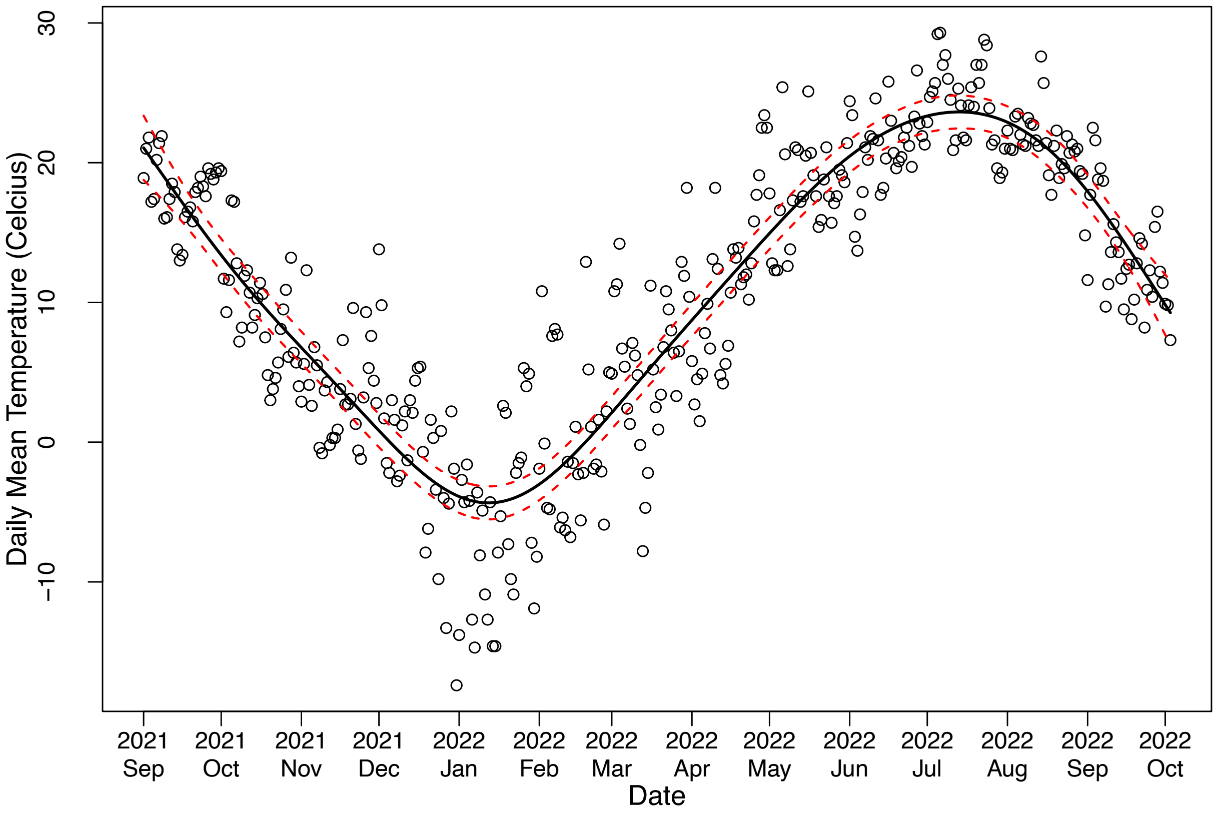


**Figure S1.** Relationship of mean daily temperature to survey date during the period when camera trapping and visual surveys were conducted for eastern gray squirrels (*Sciurus carolinensis*). Best-fit line (solid black line) and 95% confidence intervals (dotted red lines) from a generalized additive model ($R^2 = 0.83$).

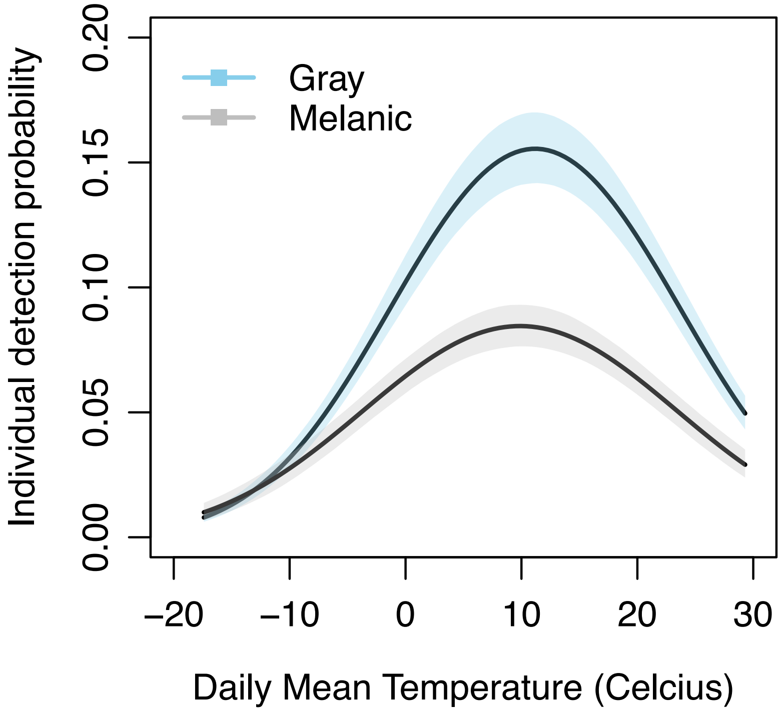


**Figure S2.** Relationship of individual detection probability of melanic and gray morphs of eastern gray squirrels (*Sciurus carolinensis*) to daily mean temperature based on a hierarchical model integrating camera trap and point count observations across N = 76 sites. Solid lines represent the predicted individual detection probabilities from the model, and the shaded areas represent 95% credible intervals.

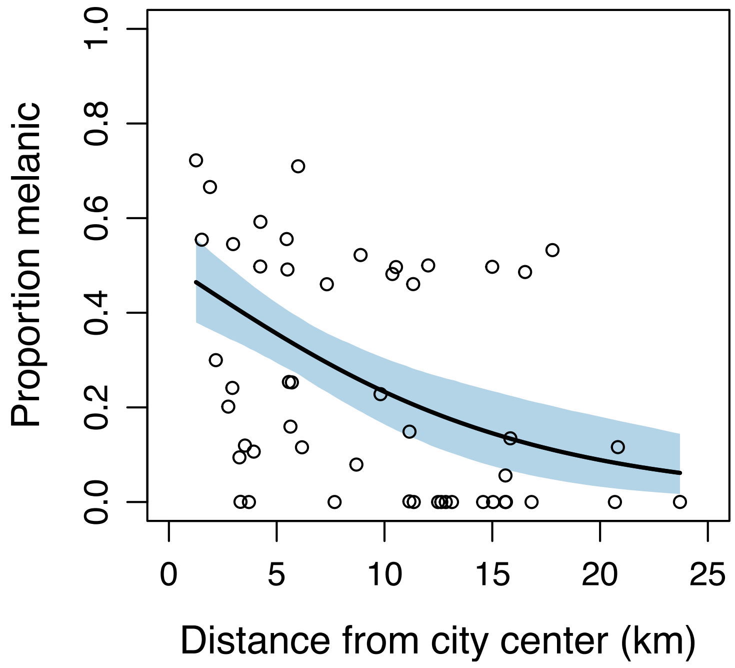


**Figure S3.** Relationship of the proportion of melanic eastern gray squirrels (*Sciurus carolinensis*) to the distance to city center for Syracuse, NY based on a hierarchical model integrating a subset of camera trap and point count observations of squirrels across N = 50 sites (16 September 2021 – 15 March 2022). Solid line represents the predicted proportion of melanic squirrels from the model, and the shaded area represents a 95% credible interval. Open circles represent estimates of the proportion melanic at each site.

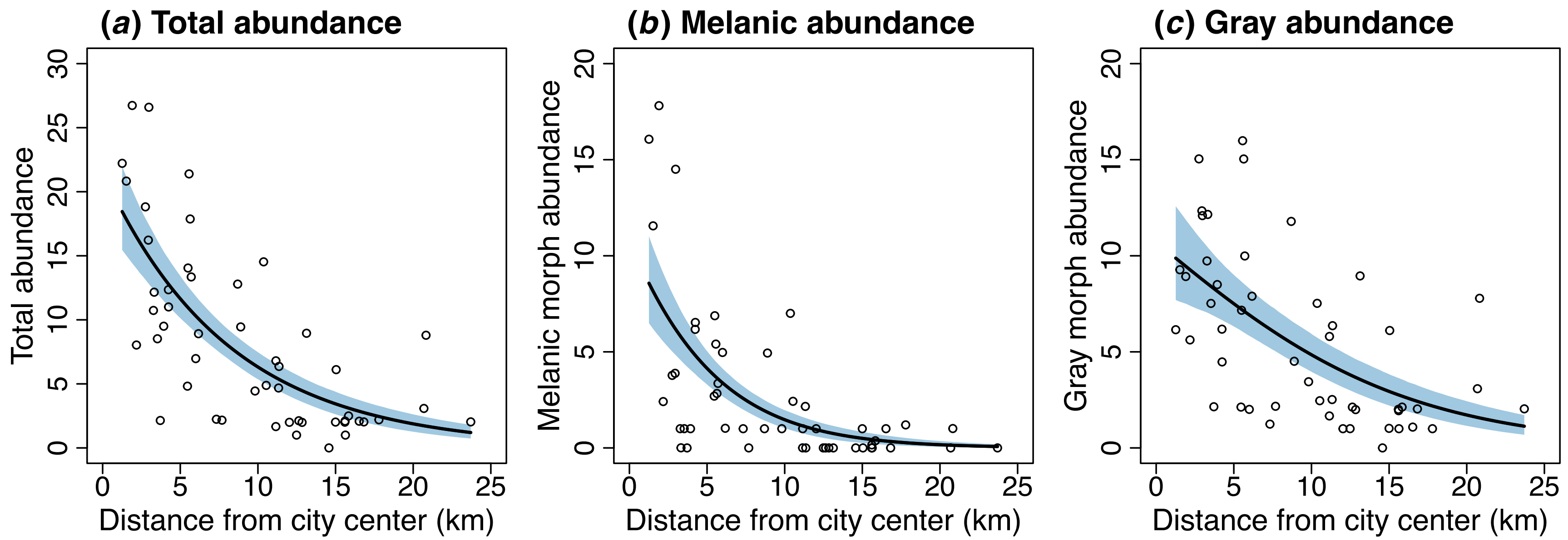


**Figure S4.** Relationship of the abundance of eastern gray squirrels (*Sciurus carolinensis*) in total (*a*) and for each color morph (*b-c*) to the distance to city center for Syracuse, NY based on a hierarchical model integrating a subset camera trap and point count observations of squirrels across N = 50 sites (16 September 2021 – 15 March 2022). Solid lines represent predicted abundance from the model, and shaded areas represent 95% credible intervals. Open circles represent estimates of abundance at each site.

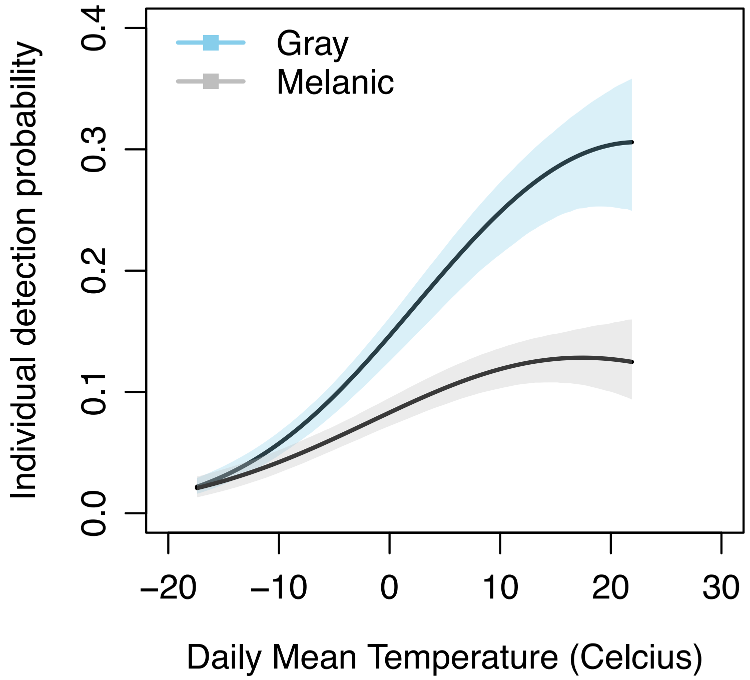


**Figure S5.** Relationship of individual detection probability of melanic and gray morphs of eastern gray squirrels (*Sciurus carolinensis*) to daily mean temperature based on a hierarchical model integrating a subset of camera trap and point count observations across N = 50 sites (16 September 2021 – 15 March 2022). Solid lines represent the predicted individual detection probabilities from the model, and the shaded areas represents 95% credible intervals.

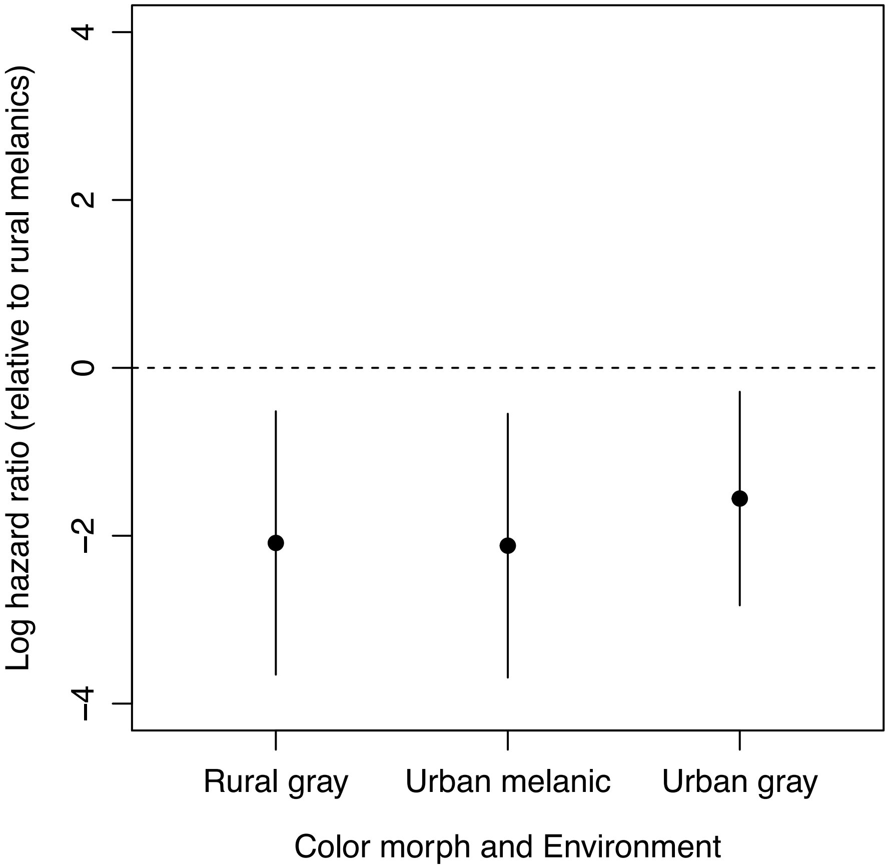


**Figure S6.** Log hazard ratios predicted from the Cox proportional hazards regression model including effects of morph, environment, and morph*environment interaction (Table 2). Log hazard ratios are shown for the rural gray, urban melanic, and urban gray morphs relative to the rural melanic morph. Error bars represent 95% confidence intervals. Hazard ratios

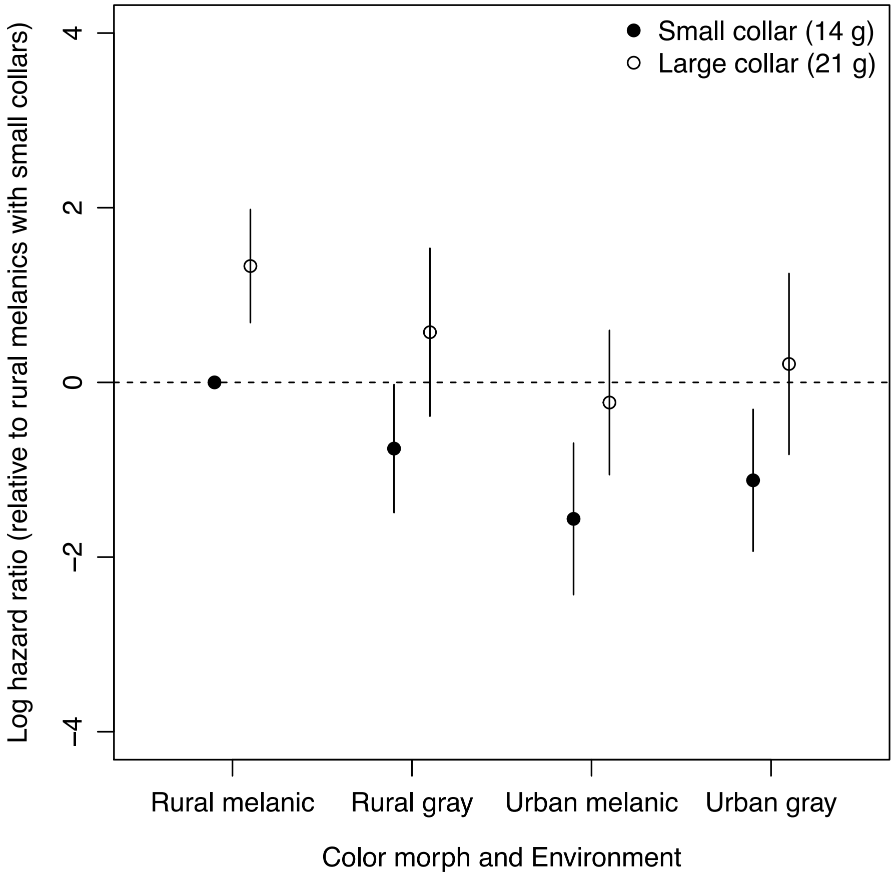


**Figure S7** Log hazard ratios predicted from the Cox proportional hazards regression model when including confirmed, probable, and possible mortalities as true mortalities. The model included effects of morph, environment, morph*environment interaction, and color size (Table S3). Hazard ratios are shown for each color morph, environment, and collar size relative to the rural melanic morph with small collars. Error bars represent 95% confidence intervals.

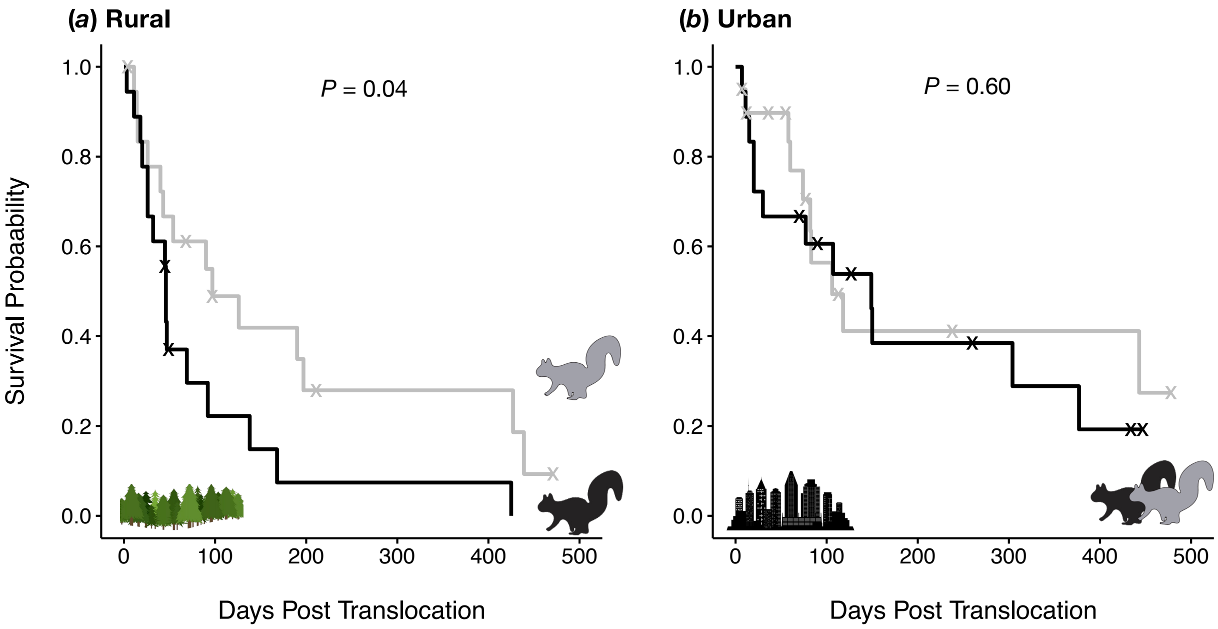


**Figure S8.** Survival curves for gray (solid gray line) and melanic (solid black line) color morphs of eastern gray squirrels (*Sciurus carolinensis*) translocated to rural (*a*) or urban (*b*) environments when including confirmed, probable, and possible mortalities as true mortalities. Steps down represent confirmed, probable, and possible mortalities, and "X"s represent missing and active squirrels that ere right-censored for the analysis. *P*-values are shown for the effect of morph within each environment based on log rank tests.

**Table S1**. Locations of sites to which eastern gray squirrels (*Sciurus carolinensis*) were translocated for survival estimation in urban and rural environments.

| Release site | Property Name | Decimal degrees (latitude, longitude) |
|---|---|---|
| Rural 1 | Three Rivers Wildlife Management Area | 43.194, -76.335 |
| Rural 2 | Otisco Lake Boat Ramp | 42.841, -76.267 |
| Rural 3 | Labrador Hollow Unique Area | 42.784, -76.057 |
| Urban 1 | Thornden Park | 43.041, -76.126 |
| Urban 2 | Westminster Park | 43.034, -76.121 |
| Urban 3 | Morningside Park | 43.028, -76.126 |
| Urban 4 | Barry Park | 43.025, -76.115 |
| Urban 5 | State University of New York College of Environmental Science and Forestry | 43.034, -76.135 |

**Table S2.** Parameter estimates for Cox proportional hazards regression model for effects of color morph (melanic, gray), environment (rural, urban), sex, body mass, collar size, and mean daily temperature during the first week of translocation on daily survival of translocated eastern gray squirrels (*Sciurus carolinensis*). Reference categories were set as melanic for color morph, rural for environment, female for sex, and small for collar size.

| Explanatory variable | Log hazard ratio | SE | *z* | *P* |
|---|---|---|---|---|
| Color morph | -3.16 | 1.14 | -2.76 | 0.006 |
| Environment | -2.78 | 1.02 | -2.74 | 0.006 |
| Color morph * Environment | 4.11 | 1.57 | 2.62 | 0.009 |
| Sex | 0.27 | 0.66 | 0.40 | 0.688 |
| Body mass | 0.00 | 0.01 | 0.18 | 0.861 |
| Collar size | 0.73 | 0.94 | 0.77 | 0.441 |
| Temperature | 2.24 | 2.91 | 0.77 | 0.441 |
| Temperature$^2$ | -0.05 | 0.06 | -0.82 | 0.413 |

**Table S3.** Parameter estimates and 95% Bayesian credible intervals (CI) for models of abundance, proportion melanic, and detection probability of eastern gray squirrels (*Sciurus carolinensis*) based on a hierarchical model integrating camera trap and point count observations across N = 76 sites. Includes estimated means, lower 95% CI bound (95% LCI), and upper 95% CI bound (95% UCI) across 3,000 iterations of the model.

| Response variable | Parameter | Mean | 95% LCI | 95% UCI |
|---|---|---|---|---|
| Total squirrel abundance | Intercept | 2.39 | 2.27 | 2.50 |
| | Distance to city center | -0.68 | -0.78 | -0.57 |
| Proportion melanic | Intercept | -1.12 | -1.39 | -0.85 |
| | Distance to city center | -0.74 | -1.02 | -0.48 |
| Melanic detection probability | Intercept | -2.39 | -2.49 | -2.28 |
| | Temperature | -0.06 | -0.11 | -0.01 |
| | $Temperature^2$ | -0.31 | -0.36 | -0.26 |
| Gray detection probability | Intercept | -1.69 | -1.80 | -1.59 |
| | Temperature | 0.03 | -0.01 | 0.06 |
| | $Temperature^2$ | -0.40 | -0.43 | -0.37 |

**Table S4.** Parameter estimates and 95% Bayesian credible intervals (CI) for models of abundance, proportion melanic, and detection probability of eastern gray squirrels (*Sciurus carolinensis*) based on a hierarchical model integrating a subset of camera trap and point count observations across N = 50 sites (16 September 2021 – 15 March 2022). A quadratic effect of survey date was included in the models for detection probability because we expected peaks in squirrel activity in fall and spring. Includes estimated means, lower 95% CI bound (95% LCI), and upper 95% CI bound (95% UCI) across 3,000 iterations of the Bayesian N-mixture model.

| Response variable | Parameter | Mean | 95% LCI | 95% UCI |
|---|---|---|---|---|
| Total squirrel abundance | Intercept | 1.90 | 1.74 | 2.06 |
| | Distance to city center | -0.71 | -0.85 | -0.58 |
| Proportion melanic | Intercept | -1.15 | -1.55 | -0.79 |
| | Distance to city center | -0.70 | -1.06 | -0.37 |
| Melanic detection probability | Intercept | -2.20 | -2.35 | -2.05 |
| | Temperature | 0.39 | 0.31 | 0.47 |
| | $Temperature^2$ | -0.14 | -0.21 | -0.07 |
| Gray detection probability | Intercept | -1.45 | -1.64 | -1.33 |
| | Temperature | 0.63 | 0.58 | 0.69 |
| | $Temperature^2$ | -0.16 | -0.21 | -0.11 |

**Table S5.** Parameter estimates for Cox proportional hazards regression model for effects of color morph (melanic, gray), environment (rural, urban), and collar size (14 g vs. 21 g) on daily survival of translocated eastern gray squirrels (*Sciurus carolinensis*) when including confirmed, probable, and possible mortalities as true mortalites. Reference categories were set as melanic for color morph, rural for environment, and small for collar size.

| Explanatory variable | Log hazard ratio | SE | *z* | *P* |
|---|---|---|---|---|
| Color morph | -0.76 | 0.37 | -2.03 | 0.043 |
| Environment | -1.56 | 0.44 | -3.53 | <0.001 |
| Color morph * Environment | 1.20 | 0.61 | 1.97 | 0.049 |
| Collar size | 1.33 | 0.33 | 4.03 | <0.001 |
